# Assessing drug uptake and response differences in 2D and 3D cellular environments using stimulated Raman scattering microscopy

**DOI:** 10.1101/2024.04.22.590622

**Authors:** Fiona Xi Xu, Rui Sun, Ryan Owens, Kailun Hu, Dan Fu

## Abstract

The architecture of cell culture—two-dimensional (2D) versus three-dimensional (3D)—significantly impacts various cellular factors, including cell-cell interactions, nutrient and oxygen gradients, metabolic activity, and gene expression profiles. This can result in different cellular responses during cancer drug treatment, with 3D-cultured cells often exhibiting higher resistance to chemotherapeutic drugs. While various genetic and proteomic analyses have been employed to investigate the underlying mechanisms of this increased resistance, complementary techniques that provide experimental evidence of spatial molecular profiling data are limited. Stimulated Raman scattering (SRS) microscopy has demonstrated its capability to measure both intracellular drug uptake and growth inhibition. In this work, we applied three-band SRS imaging to 2D and 3D cell cultures and provided a comparative analysis of drug uptake and response with the goal of understanding whether the difference in drug uptake explains the drug resistance in 3D culture compared to 2D. Our investigations revealed that despite similar intracellular drug levels in 2D and 3D A549 cells during lapatinib treatment, the growth of 3D spheroids is less impacted, supporting an enhanced drug tolerance in the 3D microenvironment. We further elucidated drug penetration patterns and the resulting heterogeneous cellular responses across different spheroid layers. Additionally, we investigated the role of the extracellular matrix in modulating drug delivery and cell response, and we discovered that limited drug penetration in 3D could also contribute to lower drug response. Our study provides valuable insights into the intricate mechanisms of increased drug resistance in 3D tumor models during cancer drug treatments.

## Introduction

Two-dimensional (2D) monolayer cell culture has traditionally been used for biological studies and drug screenings in vitro due to its simplicity and cost-effectiveness. However, 2D cell cultures are insufficient for capturing the in vivo tumor response during drug treatment, and three-dimensional (3D) cell cultures offer more intricate and representative models.^1^ Analogous to in vivo tissue such as tumors, proliferating, quiescent, and necrotic cells coexist within 3D spheroids.^2,3^ Additionally, spheroids have been reported to have similar growth dynamics to solid tumors.^4,5^ As a result, in recent years, 3D cell cultures have emerged as increasingly popular models for drug screening and cancer treatment.^6^ The more complex cell-cell and cell-matrix interactions and nutrient and oxygen gradients in 3D microenvironments result in distinct cellular structures, metabolic activities, and gene expression profiles from 2D cells.^7–11^ The differences between 2D and 3D cells can lead to varied cellular responses to drug treatments; specifically, it has been reported that 3D-cultured cells generally exhibit higher resistance to drugs.^2^

The molecular mechanisms underlying the increased resistance in 3D cells are not fully understood, yet several hypotheses have been proposed: the extracellular matrix and cell-cell interactions within the 3D microenvironment may create physical barriers that impede drug penetration^12,13^; nutrient and oxygen availability disparities may alter cellular metabolism and survival pathways^2,14,15^; different gene expression and activation of signaling pathways may promote cell survival and drug efflux^7,9,16^; and the higher cell heterogeneity and microenvironmental gradients may lead to a selection of more resistant cell populations^17,18^ Various techniques, notably genetic, transcriptomic, and proteomic analyses, have been employed to dissect these complex mechanisms. These methods, including gene expression analysis, Western blot, and multi-omics approaches, collectively identified diverse gene expressions, signaling pathways, and metabolic processes, including EGFR and EMT-associated protein expression, HER-family drug target expression, and AKT–mTORC1 signaling, that are significantly different between 2D and 3D cells and are potentially associated with reduced drug efficacy in 3D.^7,8,19–26^ While these techniques deliver essential insights into the potential causes of drug resistance evolvement, intracellular drug exposure within the complex 3D culture models, a key factor that determines heterogeneous cell response, has rarely been determined with existing techniques. Thus, it is crucial to integrate information gained from these methods with techniques that provide spatially resolved molecular profiling.

Existing techniques for directly measuring intracellular drug concentration in 3D environments are limited. Current studies largely depend on biochemical assays, which provide bulk measurements that may overlook variability in drug uptake between individual cells and their complex spatial interactions within spheroids.^27–29^ Mass spectrometry imaging has also been utilized to map drug distribution in spheroids and tissues.^30–33^ However, the method requires extensive sample preparation and cryosectioning, limiting its applications for live-spheroid imaging and cellular dynamic studies. The lack of precise 3D drug quantification could significantly hinder our understanding of in vivo drug efficacy. Typically, drug exposure within a tumor is assumed to be equivalent to plasma concentration; however, uneven or reduced drug exposure can lead to decreased treatment efficacy and the emergence of drug resistance.^34–36^ Moreover, correlating drug concentration measurement with cellular response inhibition can be highly beneficial for understanding drug-cell interactions and resistance development mechanisms.

Recently, stimulated Raman scattering (SRS) microscopy has shown great potential in both directly quantifying intracellular drug uptake and measuring single-cell growth rate as an indicator of drug response.^37,38^ In particular, Wong et al. leveraged the well-established lysosomotropic effect, in which the acidic environment in cell lysosomes protonates the weakly basic drugs and enriches drug molecules, and performed live cell hyperspectral SRS imaging in the fingerprint region to visualize and quantify drug uptakes in 2D-cultured cells.^37^ However, the compatibility of this technique with 3D cells has not been explored. Due to light scattering, the quantification of drug concentration becomes a challenge. Meanwhile, Xu et al. employed deuterated amino acids, leucine, isoleucine, and valine (d-LIV), as tracers and measured the ratio of carbon–deuterium (C– D) SRS signal from deuterium-labeled proteins and the carbon–hydrogen (C–H) SRS signal from unlabeled proteins to determine the cell growth rate of 2D and 3D cells.^38^ However, this method has only been applied to fixed cells.

In this work, we present a robust, noninvasive three-band SRS imaging method that integrates the measurements of drug uptake and cellular response of live cells across 2D and 3D microenvironments. We report the drug uptake and response differences between cells in 2D and 3D during drug treatments and closely examine the heterogeneous drug penetration and cell response patterns within 3D spheroids. Furthermore, by comparing different 3D spheroid culture methods, we explore the effects of extracellular matrix on drug delivery and cell growth inhibition. Overall, this work introduces a valuable tool for innovative experimental measurements of drug uptake and response differences between 2D and 3D cellular environments. It illuminates the intrinsic differences in cellular response to drugs under different microenvironments and the contribution of limited drug penetration in 3D to drug resistance. Our findings offer empirical evidence to understand the complex mechanisms of drug resistance development in 3D cell models and have important implications in drug screening assay development as well as cancer therapies.

## Materials and Methods

### Three-band SRS imaging

The hyperspectral SRS microscopy setup has been reported in detail in our previous work.^37^ Briefly, a broadband femtosecond dual beam laser system (Spectra-Physics Insight DS+) outputs a wavelength-tunable beam (pump) and a fixed beam at 1040 nm (Stokes) at an 80 MHz repetition rate. The pump beam is chirped to approximately 3 ps with high-dispersion SF11 dense flint glass rods (GR, Newlight Photonics). The Stokes beam is modulated at 20 MHz with an electro-optical modulator (EOM, Thorlabs) and coupled into a 4-m polarization-maintaining Yb-doped fiber (YB1200-10/125 DC-PM, Thorlabs) for parabolic amplification^39^, then chirped to approximately 3 ps using a grating stretcher (GS, LightSmyth). The two beams are combined with a 1000 nm short pass (SP) dichroic mirror (Thorlabs) and temporally overlapped using a motorized delay stage (DS, Zaber X-DMQ-AE).

The synchronized laser beams are directed into a home-built upright laser scanning microscope equipped with a 40× water immersion objective (Nikon N40XLWD-NIR, NA = 1.15) and an oil immersion condenser (Nikon MEL41410, NA = 1.4). The pump beam is isolated with a 1000 nm SP filter (Thorlabs) and detected by an amplified Si photodiode (PD). The signal is then demodulated with a lock-in amplifier (LIA, Zurich Instruments H2FLI) with a time constant of 4 μs to generate 512 × 512-pixel SRS images. For SRS imaging of live cells and spheroids, 40 mW of the pump beam at 800 nm for C–H, 858 nm for C–D, and 912 nm for fingerprint region, and 90 mW of the Stokes beam were used.

### 2D cell culture

The human lung adenocarcinoma cell line, A549 cells (ATCC), was maintained at 37 °C in a humidified 5% CO_2_ incubator and cultured in Dulbecco’s modified Eagle’s medium (DMEM, Gibco) supplemented with 10% fetal bovine serum (Hyclone) and 1% penicillin-streptomycin (Gibco). Cells were seeded at 200,000 cells per coverslip and incubated for 48 hours before drug treatment and deuterium labeling.

### 3D spheroid culture

For scaffold-free spheroid culture, the A549 cell line cultured in DMEM supplemented with 10% FBS and 1% penicillin-streptomycin prior to the 3D spheroid culture was washed twice with PBS and detached from the dish with 0.25% trypsin (Gibco), then centrifuged at 200 × g for 5 min at room temperature and resuspended in medium. The cellular suspension was diluted and transferred into each well of the 96-well low-attachment culture plate (Corning) to achieve 2000 cells/well. The culture plate was centrifuged at 200 × g for 3 min at room temperature and incubated at 37 °C in a humidified 5% CO_2_ incubator for 48 hours before drug treatment and deuterium labeling.

For scaffolded extracellular matrix spheroid culture, A549 cellular suspension was counted and diluted to 400,000 cells/mL. Matrigel (Corning) was stored under −20°C and thawed under 4°C overnight prior to use. A one-to-one ratio of Matrigel and diluted cellular suspension was mixed, and 35 μL of the mixture was transferred onto the center of a coverslip. The Matrigel and cell mixture was prepared on an ice block. The coverslip was then inverted so that the Matrigel and cell mixture was at a hanging drop orientation and incubated in a 37°C humidified 5% CO_2_ incubator for 30 mins to allow polymerization. Then the coverslip was flipped back, and the Matrigel droplet was cultured with the immersion of DMEM in the incubator for 2-3 weeks before drug treatment and deuterium labeling, during which the culture medium was replaced every 2 days.

### Deuterium labeling for single-cell growth rate measurement

Deuterated L-leucine-d_10_, L-isoleucine-d_10_, and L-valine-d_8_ (d-LIV, Cambridge Isotope Laboratories) were supplemented into the LIV-deficient DMEM medium (Boca Scientific) at a concentration of 0.8 mM. This concentration matches the concentrations of these amino acids in regular DMEM to achieve full deuteration. To perform deuterium labeling, the cell culture medium was changed from the regular DMEM to the d-LIV-supplemented medium, followed by incubation at 37 °C for the desired labeling time. Specifically, 2D-cultured cells were labeled with d-LIV for 6 hours before SRS imaging, and 3D-cultured spheroids were d-LIV labeled for 24 hours.

### Single-cell and single-spheroid growth rate quantification

Our previous work has reported details on measuring single-cell growth rates with ratiometric SRS imaging.^38^ Briefly, d-LIV-labeled cells were imaged at 2125 cm^−1^ in the C–D region to quantify their newly synthesized deuterium-labeled proteins and at 2930 cm^−1^ in the C– H stretching region to measure their total unlabeled proteins. In addition, SRS images at 2030 cm^−1^ were collected as the off-resonance frequency to remove the non-Raman background signal in the C–D SRS images. For 2D-cultured cells, we generated cell masks by thresholding the C–H SRS image and segmenting adjacent cells with the watershed algorithm. For 3D cells, each spheroid was manually segmented using its C–H SRS image. The cell and spheroid masks were then used to calculate C–D/C–H SRS ratios as growth rate measurements.

### Drug treatment for intracellular drug uptake measurement

Lapatinib (Selleckchem), afatinib (Selleckchem), osimertinib (Selleckchem), and etoposide (Sigma-Aldrich) were dissolved in dimethyl sulfoxide (DMSO) to make 10 mg/mL stock solutions. The stock solutions were diluted with cell culture medium to the desired concentrations for drug treatment. Control cells were treated with DMSO only. Drug solutions were prepared using the d-LIV-supplemented medium for deuterium labeling. For both 2D and 3D cells, the drug treatment time was 24 hours.

### Intracellular drug uptake quantification

Details on measuring single-cell drug uptake with fingerprint region hyperspectral SRS imaging have been previously reported.^37^ Briefly, live A549 cells and spheroids treated with lapatinib were imaged at the 1300 – 1600 cm^−1^ spectral band, which captures the cell peak at 1450 cm^−1^ and lapatinib peaks. The single-cell or single-spheroid SRS spectrum was extracted with cell or spheroid masks generated from C–H SRS images as described above. A spectral unmixing algorithm was then performed using the lsqr function in MATLAB to determine the linear combination coefficients of cell, lipid droplet, drug, and background solvents. In the previous work, the absolute concentration was determined by comparing the drug signal to calibration. In the presence of light scattering in 3D spheroids, the intracellular drug uptake was quantified as the calculated coefficient of the drug component divided by its cell signal intensity at 1450 cm^−1^. This normalization step is necessary to remove the contribution of light scattering-induced intensity loss.

## Results and Discussion

We first validated the capability of measuring both intracellular drug uptake and growth rate inhibition of the same cells in 2D using our three-band SRS imaging system. We selected A549 cells, a non-small-cell lung cancer cell line, as the cell model for this study. We treated the cells with lapatinib (Lap), an FDA-approved tyrosine kinase inhibitor (TKI) for both the epidermal growth factor receptor (EGFR) and human epidermal growth factor receptor 2 (HER2), at varying concentrations for 24 hours. The SRS images in three different vibrational bands, fingerprint, C– D, and C–H regions, were acquired for 150-300 cells for each treatment. We performed hyperspectral SRS imaging in the 1300 – 1600 cm^−1^ spectral band to quantify the single-cell drug uptake. Representative SRS images and spectra of cells treated with 5 μM of lapatinib were shown in **Figure 1A-B**. Lapatinib has 4 distinct peaks, at 1325, 1360, 1390, and 1535 cm^−1^, that are well-separatable from the 1450 cm^−1^ cell peak in this region. We detected strong drug signals at its most prominent peak in the imaged spectral region, at 1360 cm^−1^ (**Figure 1A** left), consistent with previous reports that the drug is highly enriched in lysosomes to reach a detectable range.^37,40^ However, the drug concentration is not quantifiable from this single SRS image as signals from cells also contribute to this frequency (**Figure 1B**). Leveraging the linear dependence of the SRS signal on chemical concentration, we apply a linear least-squares algorithm on cell spectra to unmix the cellular and drug compositions. To correct for signal variation induced by light scattering, we quantify single-cell intracellular drug uptake by normalizing the drug coefficient to the 1450 cm^−1^ cell peak, resulting from CH_2_ deformation of protein and lipid. We plotted 2D cell drug uptake as a function of lapatinib concentrations in **Figure 1C**. The 2D cells were dosed with up to 10 μM lapatinib only, as higher drug concentrations dramatically diminished cell viability. Overall, as the drug dosage increases, the measured intracellular drug uptake increases as expected, confirming the sensitivity of our method. To confirm that the normalization by 1450 cm^−1^ does not bias results, we also normalized the drug coefficient to the C–H stretching peak at 2930 cm^−1^ and obtained the same trend, as demonstrated in **Figure S1 A**. Although both cell peaks at 1450 cm^−1^ and 2930 cm^−1^ can be used for the normalization, light scattering is wavelength-dependent, and therefore, it is more desirable to use the 1450 cm^−1^ peak despite its higher noise since the same wavelength is used for excitation of the peaks in the fingerprint region.

**Figure 1.**
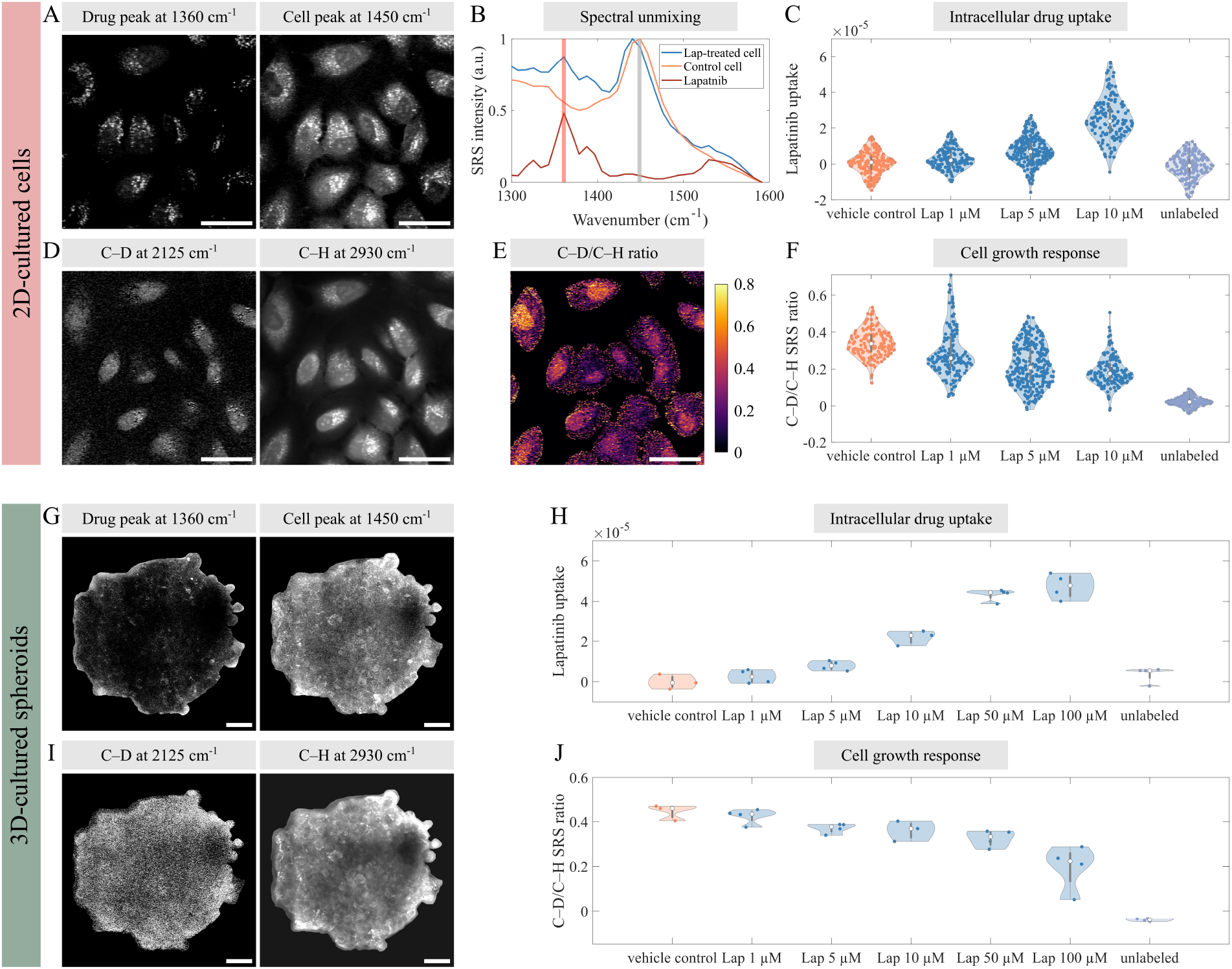
Representative SRS images and spectra of live 2D and 3D A549 cells and their intracellular drug uptake and growth rate changes during treatment with a tyrosine kinase inhibitor, lapatinib (Lap). (A) SRS image of 2D A549 cells treated with 5 μM of lapatinib at 1360 cm^−1^, the most prominent drug peak (red shaded line in B), and at 1450 cm^−1^, the cell peak (gray shaded line in B). (B) SRS spectra of Lap-treated cells were decomposed into cellular and drug components using a linear least-squares algorithm to measure drug uptakes. (C) The intracellular drug uptake in 2D cells treated with Lap at varying concentrations for 24 hours. (D) SRS images of 2D A549 cells at 2125 cm^−1^ in the C–D region and 2930 cm^−1^ in the C–H region. Their ratiometric image (E) reflects cellular growth rate. (F) The growth rate response of 2D cells labeled with d-LIV for 6 hours under different lapatinib treatments. (G, I) SRS image of 3D spheroid treated with 5 μM of lapatinib at drug, cell, C–D, and C–H peaks. (H, J) Single-spheroid drug uptake and growth rate response under lapatinib treatment of varying concentrations and 24-hour d-LIV labeling. Scale bars: 40 μm.

To measure the single-cell growth rate, we labeled the 2D A549 cells with d-LIV for 6 hours, which provides a sufficient C–D signal-to-noise ratio (SNR) for reliable quantification (**Figure 1D**). **Figure 1E** shows the C–D/C–H ratiometric image that allows visualization of different cell growth rates. The 2D single-cell growth response was plotted as a function of lapatinib concentration in **Figure 1F**. It has been reported that lapatinib significantly reduces various cellular activities in A549 cells, including cell proliferation, DNA synthesis, and colony formation capacity.^41^ As the concentration of drug treatment increases, the C–D/C–H ratios decrease, agreeing with the expected growth inhibition.

We then performed the three-band SRS imaging method to quantify drug uptake and growth response in 3D cells. We cultured A549 cells in low-attachment plates to form spheroids and treated the spheroids with up to 100 μM lapatinib. 3-4 spheroids were imaged under each treatment. Representative drug peak, cell peak, C–D, and C–H SRS images of the middle cross-section of a spheroid treated with 5 μM of lapatinib are shown in **Figure 1G** and **I**. It has been reported that 3D-cultured cells have a slower proliferation rate compared to monolayer cells^2,9,16^; therefore, we performed 24-hour d-LIV labeling on 3D spheroids to obtain a sufficient C–D signal (**Figure 1I**). Different from 2D cell imaging, mosaic tile scans were needed to capture an entire spheroid cross-section. However, as illustrated in **Figure S2 A**, the field curvature of each field-of-view (FOV) results in obvious stitching artifacts that can greatly reduce the accuracy of our measurements. To address this problem, we generated an FOV normalization mask using cell-free regions for each spectral region. The masks were normalized to have an intensity ranging from 0 to 1. We then divided each tile image by the FOV mask to correct for the field curvature. As shown in **Figure S2 B**, this FOV normalization method significantly minimizes stitching artifacts and resultant intensity variations.

We observed stronger drug signals around the outer ring of the spheroids and weaker signals in their inner layers (**Figure 1G**). We note that cells in the center of the spheroid experienced much stronger light scattering and sample-induced aberration and thus have lower SRS signals.^42^ This is reflected in all four channel images. This problem can be addressed by normalizing the drug uptake measurement to the 1450 cm^−1^ cell peak intensity. The C–D/C–H ratio also normalizes this contribution and provides quantitative single-cell growth measurement. **Figure 1H** and **J** demonstrate the single-spheroid drug uptake and growth rate response as a function of lapatinib concentration. As the drug dosage increases, the measured intracellular drug uptake increases, and the growth rate decreases. Interestingly, the increase in intracellular drug concentration began to level off at concentrations above 50 μM. As stated above, our SRS method primarily quantifies the intracellular drug uptake through the lysosomotropic effect, and pH-driven lysosomal sequestration is saturable.^43^ The intracellular drug uptake level of spheroid treated with 50–100 μM likely indicates saturated lapatinib accumulation. These trends confirm the capability of our SRS method to detect and quantify both 2D and 3D drug uptake and cellular response simultaneously during treatments.

To further investigate differences in drug uptake and cellular response between 2D and 3D environments, we compared the intracellular lapatinib uptake, cell growth rates, and normalized cell growth rates for cells treated with the same concentrations in **Figure 2. Figure 2A** demonstrates that intracellular drug uptake levels are similar between single cells cultured in 2D and single spheroids in 3D, with no significant differences observed within each treatment concentration. The same trend was observed when using C–H peak intensity for drug uptake measurement normalization (**Figure S1 C**). To compare growth inhibition differences between 2D and 3D-cultured cells, we adjusted the C–D/C–H ratio of 2D-cultured cells by multiplying it by a factor of 4, accounting for differences in d-LIV labeling duration. As shown in **Figure 2B**, 2D cells have significantly higher growth rates compared to 3D cells under identical treatment conditions. Interestingly, when we compared the relative growth rate changes during drug treatment by normalizing 2D and 3D cell growth rates to the median growth rates of their respective vehicle controls, we found that cells in the 3D environment exhibited higher normalized growth rates than those in the 2D environment, as illustrated in **Figure 2C**. This observation confirms that, with similar levels of drug uptake, the proliferation of 3D-cultured cells is more resistant to lapatinib treatment than their 2D-cultured counterparts, consistent with our observation that 3D cell culture can tolerate much higher concentration lapatinib treatment. This trend aligns with our previous drug response study of A549 cells treated with another TKI, gefitinib, in which the drug dosage that reduces cell growth rate by half increased by ∼7-fold from 2D to 3D.^38^

**Figure 2.**
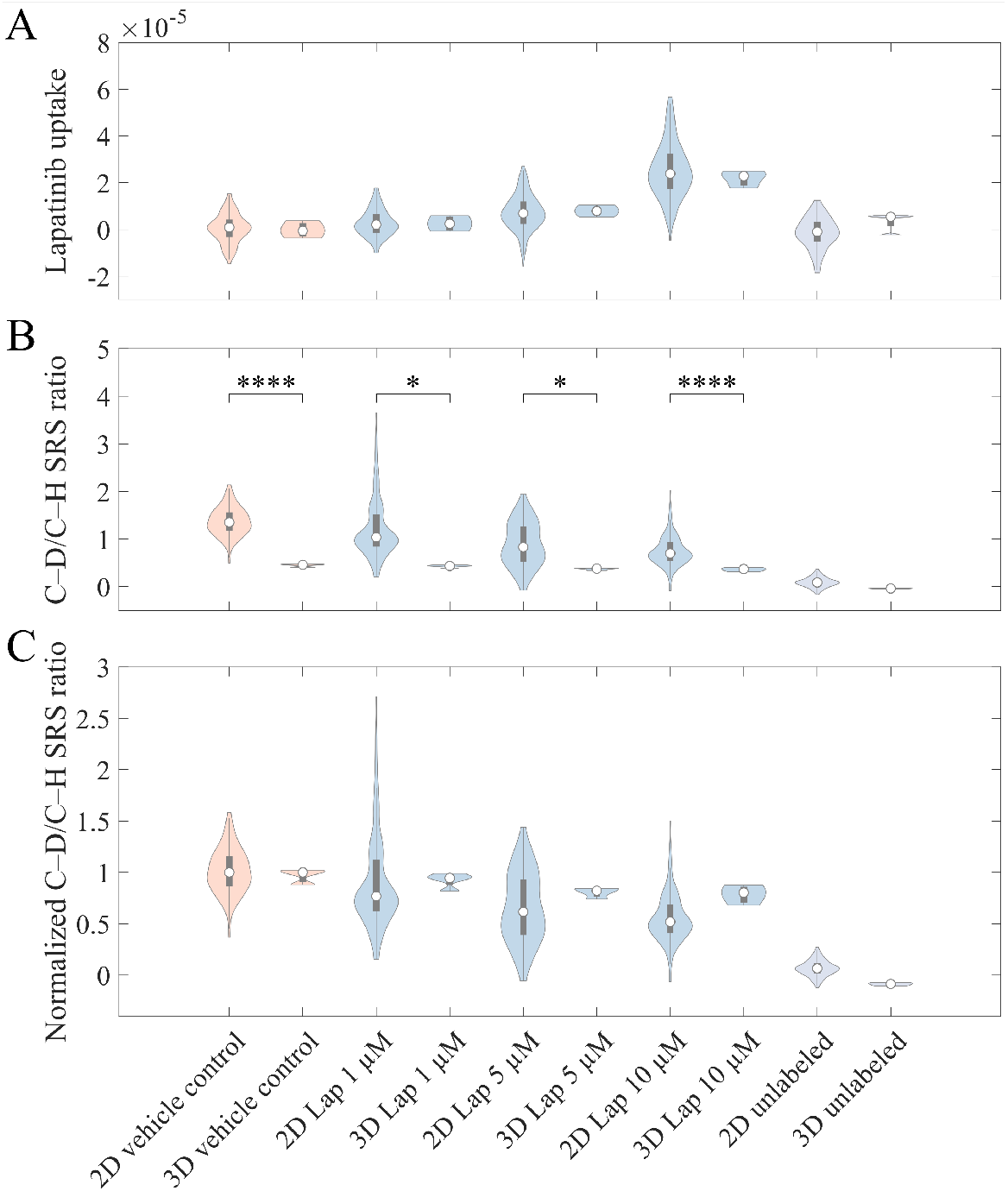
Comparison of intracellular drug uptake (A), cell growth rate (B), and normalized cell growth (C) between 2D-cultured cells and 3D-cultured spheroids treated with varying concentrations of lapatinib. Significance determined by Student’s t-test: *: p ≤ 0.05, ****: p ≤ 0.0001.

It is important to note that the 3D measurements in **Figure 2** represent the averaged data from all cells in the imaged cross-section, resulting in much smaller variances compared to 2D measurements. This averaging does not capture the heterogeneity in drug uptake or response of cells within the individual spheroid. To explore potential reasons behind the increased resistance observed in 3D-cultured cells, we examined the correlation between drug penetration and cell response heterogeneity. Given the challenge of segmenting individual cells within each spheroid due to their dense packing and the limited SNR in the spheroid center, we segmented the spheroids into layers to study how the drug diffuses and influences cell responses. Using spheroids treated with 10 μM of lapatinib as an example, we compared their drug uptake and response at different layers against control spheroids. **Figure 3A** shows the drug/C–H ratiometric SRS image of a representative Lap 10 μM-treated spheroid. Such ratiometric images enable direct visualization of drug accumulation and localization within the spheroids. **Figure 3B** provides a zoom-in view of a drug-enriched area. By comparing the ratiometric image with the C–H SRS image, we observed a high spatial correlation between lapatinib accumulation, appearing as bright yellow spots, and cell cytoplasm. We note that the drug uptake measurements were normalized to the 1450 cm^−1^ cell peak. The C–H image was used here only for this visualization because of its higher SNR. Overall, we observed significant aggregations of lapatinib molecules in the outer and middle layers of the spheroid. This observation matches the lapatinib uptake plot in **Figure 3C**. The intracellular drug concentration is at a constant level between the boundary of the spheroid and around 60 μm from the center, then decreases rapidly as getting closer to the spheroid center.

**Figure 3.**
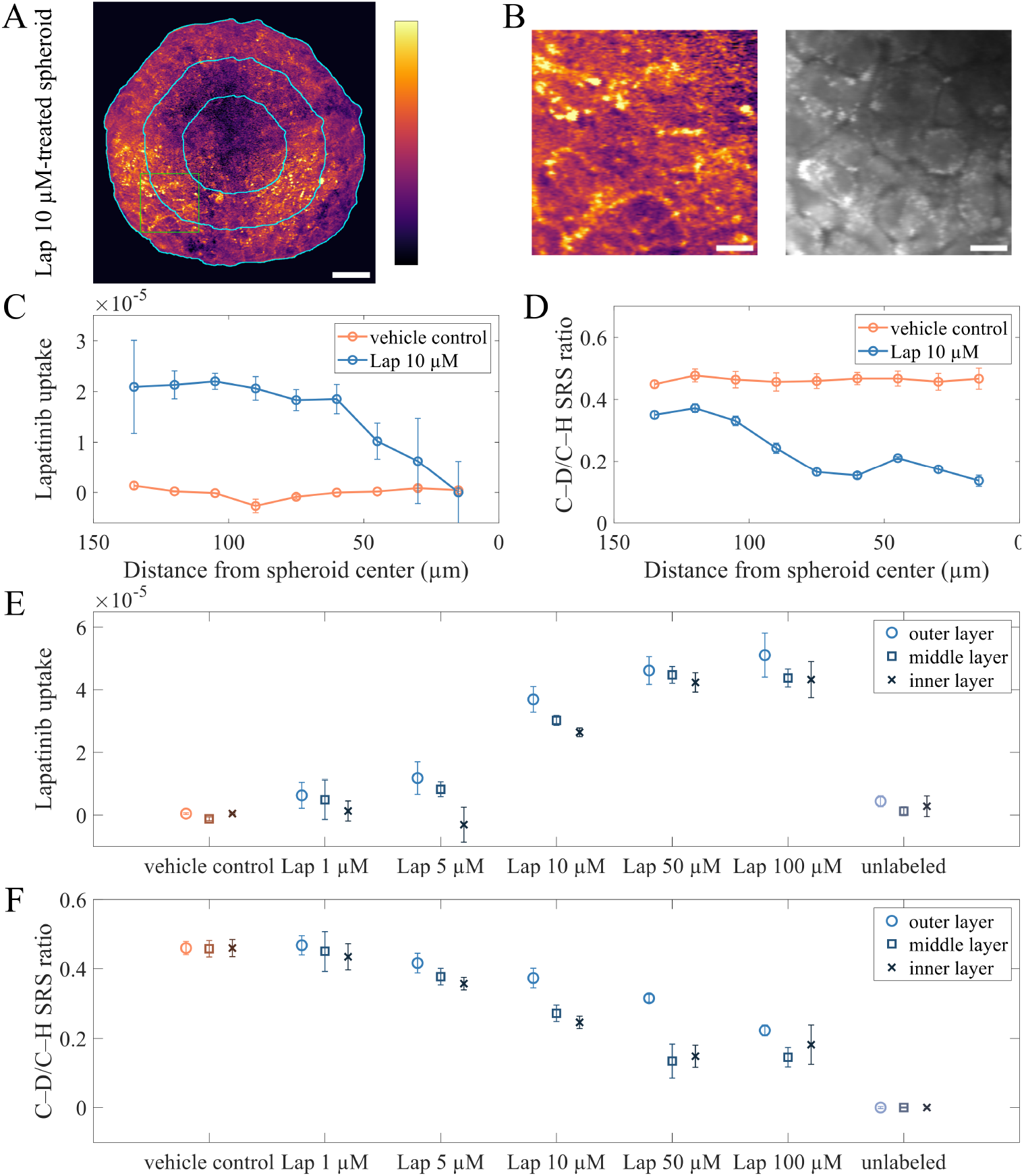
Cellular drug uptake and response in 3D-cultured spheroids treated with lapatinib. (A) The drug/C–H ratiometric image of a representative spheroid treated with 10 μM of lapatinib. (B) zoom-in ratiometric image and the corresponding C–H SRS image of a drug-enriched area (region indicated by green square in A). Spheroids were separated into layers to investigate drug penetration and cell response heterogeneity. (C-D) Plots of intracellular lapatinib uptake (C) and growth rate (D) of cells at different distances from the center of control and Lap-treated spheroids. (E-F) Plots of intracellular drug uptake (E) and growth rate (F) of cells in the inner, middle, and outer layers of spheroids treated with different concentrations of lapatinib (3 layers indicated by cyan circles in A). Scale bars: (A) 40 μm; (B) 10 μm.

**Figure 3D** illustrates the growth rates of cells at different layers in the spheroids. Cells in the vehicle control spheroids exhibited similar growth rates across all layers. It is important to note that the spheroids studied in this work have diameters of less than 300 μm and, therefore, do not have a significant oxygen concentration and nutrient penetration gradient. Spheroids exceeding 400–500 μm in diameter begin to develop a necrotic core^44^, where we would expect a reduced cell growth rate at the center. Cells in all layers of the spheroids treated with 10 μM of lapatinib exhibited decreased growth rates. Interestingly, despite having a significantly lower intracellular drug uptake, cells closer to the spheroid’s center exhibited a greater reduction in growth rates. This finding is counterintuitive, and we hypothesize that inner layer cells are potentially more sensitive to drugs than outer cells. It is possible that inner layer cells, having more limited access to nutrients and oxygen, are more likely to enter a quiescent state under treatment, in which cells are not proliferating or responsive to the drugs.^45–47^ It has been studied that quiescent cells can lead to the emergence of therapy resistance and tumor recurrence^48,49^, thus suggesting that the drug-induced quiescence of these cells could paradoxically enhance their survival capabilities and eventually contribute to increased resistance. However, further investigations are needed to elucidate the actual reasons behind the observed phenomenon.

We observed similar trends in spheroids treated with different drug concentrations. The imaged spheroids were divided into three layers, inner, middle, and outer, as illustrated in **Figure 3A** with cyan circles. **Figure 3E** displays lapatinib uptake in cells across three spheroid layers under varying dosages. With a lapatinib dosage of 1–50 μM, we can see a penetration gradient where the intracellular drug uptake level is the highest at the outer layer and lowest in the inner layer. When the drug dosage is below 5 μM, cells in the inner layer have no observable drug uptake. As dosage increases, the intracellular drug uptake level in all layers increases. As intracellular lapatinib uptake saturates in outer layer cells, drug uptake in inner cells increases until a similar level is reached across all layers. These results indicate that the drug diffusion and penetration gradient across spheroids are concentration-dependent. **Figure 3F** shows the corresponding cellular responses. Overall, as lapatinib concentration increases, the growth rates of cells in all layers decrease. Similar to the trend we discussed above, cells in the inner layers exhibit higher growth inhibition. Such an effect was more obvious at higher drug concentrations. By combining drug uptake and growth rate measurements from the same spheroids, we gained valuable insights into how drugs of different concentrations penetrate through 3D-cultured cells and consequently lead to heterogeneous cellular response gradients.

Next, we explored the effects of the extracellular matrix (ECM) on drug uptake and response of 3D spheroids. The spheroids studied in the previous sections were cultured with low-attachment plates, which is a scaffold-free approach. Although scaffold-free 3D culture methods have many advantages, including high throughput and better reproducibility, they fail to provide cell-matrix interactions in tissue. Here, we used Matrigel, a protein-based hydrogel that is a well-established biomedical material for growing spheroids.^50,51^ **Figure 4A** shows the C–H SRS images depicting a selection of representative spheroids cultured in Matrigel, showcasing significant variations in their size, shape, and structure. Most of the spheroids we selected for this study range from 200 to 400 μm in diameter. Compared to the round and densely packed spheroids observed in the scaffold-free method, as shown in **Figure 1** and **Figure 3**, scaffold-based spheroids exhibit a more irregular shape and morphology. Notably, scaffold-based spheroids often display distinct cell-free regions within their structures, a characteristic not observed in scaffold-free spheroids. We postulate that these differences are likely caused by the distinct spheroid formation processes between the two approaches. In scaffold-free methods, spheroids are formed solely through cell aggregation without the presence of surrounding supporting materials. Cells aggregate and adhere to each other, facilitated by cell-cell adhesion molecules, resulting in a tightly packed spherical structure.^52–54^ Subsequent cell growth and division lead to the enlargement of these spheroids. Conversely, in scaffold-based methods such as Matrigel culture, cells initially adhere to the surrounding extracellular matrix, forming an outer layer of the spheroid. Over time, these cells proliferate and gradually fill the inner regions of the spheroids, resulting in holes of various sizes and shapes between cells.

**Figure 4.**
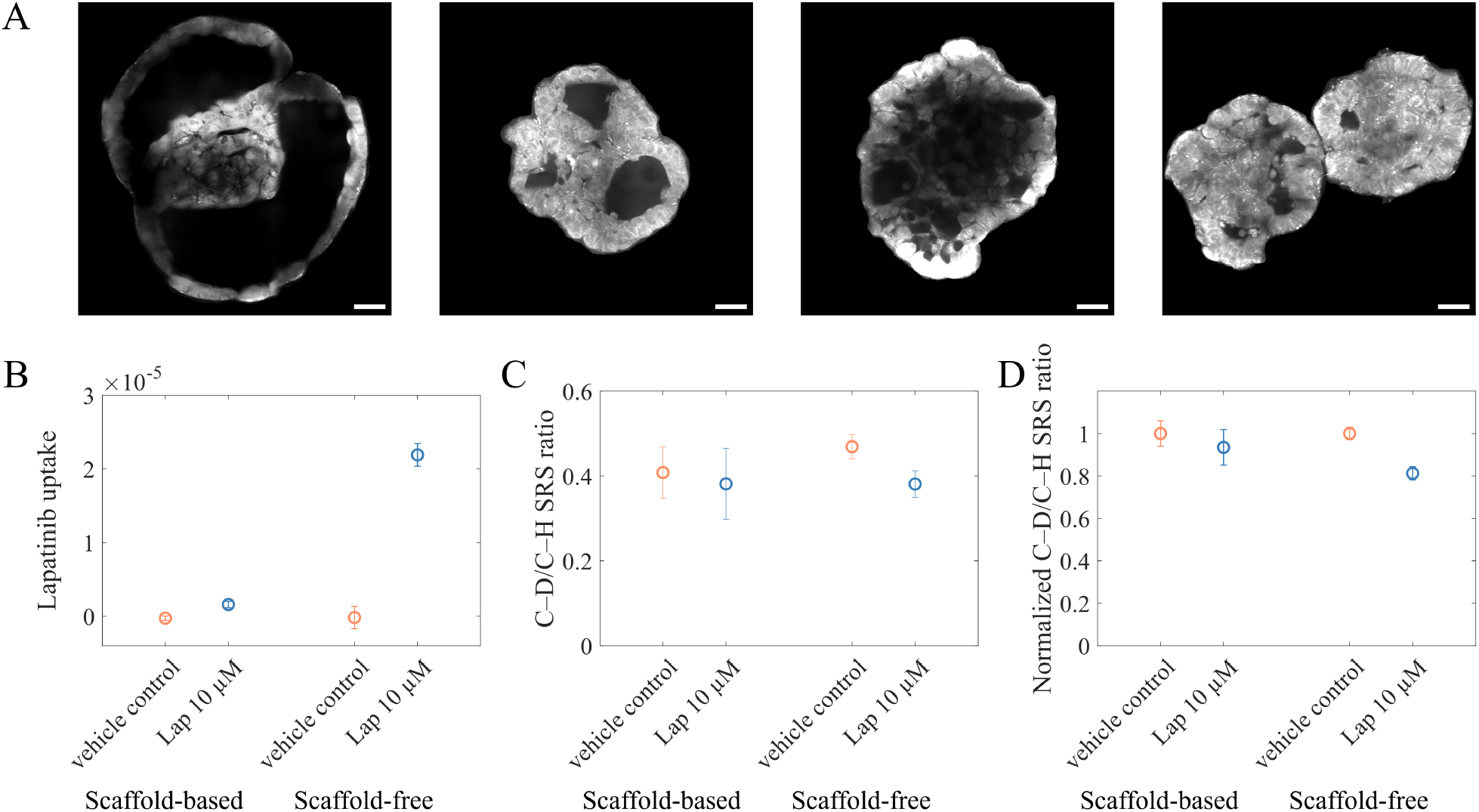
Effects of extracellular matrix on lapatinib uptake and response of 3D cells. (A) Example C–H SRS images of 3D spheroids cultured in Matrigel. (B-D) Intracellular drug uptake (B), growth rate (C), and normalized growth rate (D) comparison between scaffold-based and scaffold-free 3D spheroids treated with lapatinib at 10 μM and d-LIV-labeled for 24 hours. Scale bars: 40 μM.

To study the impact of the ECM on drug delivery and penetration, we treated Matrigel-cultured spheroids with 10 μM of lapatinib for 24 hours. We then compared the lapatinib uptake in these spheroids to that in scaffold-free spheroids under the identical treatment. The single-spheroid drug uptake measurements were normalized by the 1450 cm^−1^ cell peaks. **Figure 4B** demonstrates that the drug uptake level in scaffold-based spheroids is significantly lower than in scaffold-free spheroids. Contrary to scaffold-free culture, where spheroids are directly exposed to drug solutions, the presence of the ECM in scaffold-based culture creates a physical barrier for drug diffusion. The observed disparity in drug uptake levels between the two culture methods highlights the significant role of the ECM in impeding drug penetration in the 3D tumor environment.

Following our investigation into drug delivery and uptake, we compared spheroid growth rates between the two culture methods, as shown in **Figure 4C**. The control Matrigel-cultured spheroids exhibited slightly lower C–D/C–H SRS ratios, indicating a variance in growth rates. This reduction in growth rate can be attributed to the presence of ECM, which sterically hinders cell motion and migration and causes a less efficient nutrient delivery.^9^ Additionally, both control and drug-treated scaffolded spheroids displayed a larger variation in their growth rates compared to scaffold-free spheroids, likely arising from their diverse sizes, shapes, and structures. Furthermore, by normalizing the growth rates of drug-treated spheroids to those of their respective controls, we observed a smaller decrease in growth rates for scaffolded spheroids, depicted in **Figure 4D**. This is likely due to their significantly lower drug exposure, underscoring how the extracellular matrix influences not only drug penetration but also the resultant cellular responses to treatment. Notably, contrasting the significant reduction in drug uptake due to the extracellular matrix, the inhibition of spheroid growth was less pronounced. This suggests that while spheroid growth inhibition is related to drug concentration, the relationship is not linear. Given the much higher drug tolerance of 3D spheroids, modest growth rate inhibition changes at 10 μM of lapatinib treatment are expected despite large drug uptake changes. Overall, our findings highlight the profound impact of spheroid culture conditions on drug delivery and cellular responses, demonstrating that the physical and biochemical environment significantly modifies the effectiveness of therapeutic interventions.

Lastly, we investigated differences in 2D and 3D cell growth rate response to a chemotherapeutic drug, paclitaxel (Pac). We first calculated the drug concentration at which the 2D cell growth rate is reduced by half (GR_50_) using fixed A549 cells. The cells were treated with paclitaxel at varying concentrations, ranging from 0.2 to 5000 nM, for 24 hours. As shown in **Figure 5A**, by applying a sigmoidal fitting on the growth rate inhibition as a function of paclitaxel concentration, the GR_50_ concentration of 2D cells was determined to be 47.5 nM. Based on this measurement, we dosed cells and spheroids with 10, 50, and 100 nM of paclitaxel and performed live-cell SRS imaging after 24 hours of treatment. Representative C–H SRS images of drug-treated 2D and 3D cells are shown in **Figure 5B**. In the 2D cellular environment, at a concentration of 10 nM, many cells showed morphological features corresponding to mitotic arrest and became multinucleate. When the dosage is as high as 100 nM, most 2D A549 cells lose their nuclei and other cellular contents, indicating mitotic cell death. These observations align with the reported treatment outcomes of paclitaxel as a microtubule-stabilising drug.^55^ Spheroids exhibited much less change under the same treatments. At a dosage of 10 nM, only very few cells around the spheroid periphery rounded, while the majority of cells maintained a healthy morphology. At a paclitaxel dosage of 100 nM, cells around the spheroid periphery rounded up, and their nuclei disappeared, showing signs of mitotic arrest. However, most cells in the spheroid remained less affected.

**Figure 5.**
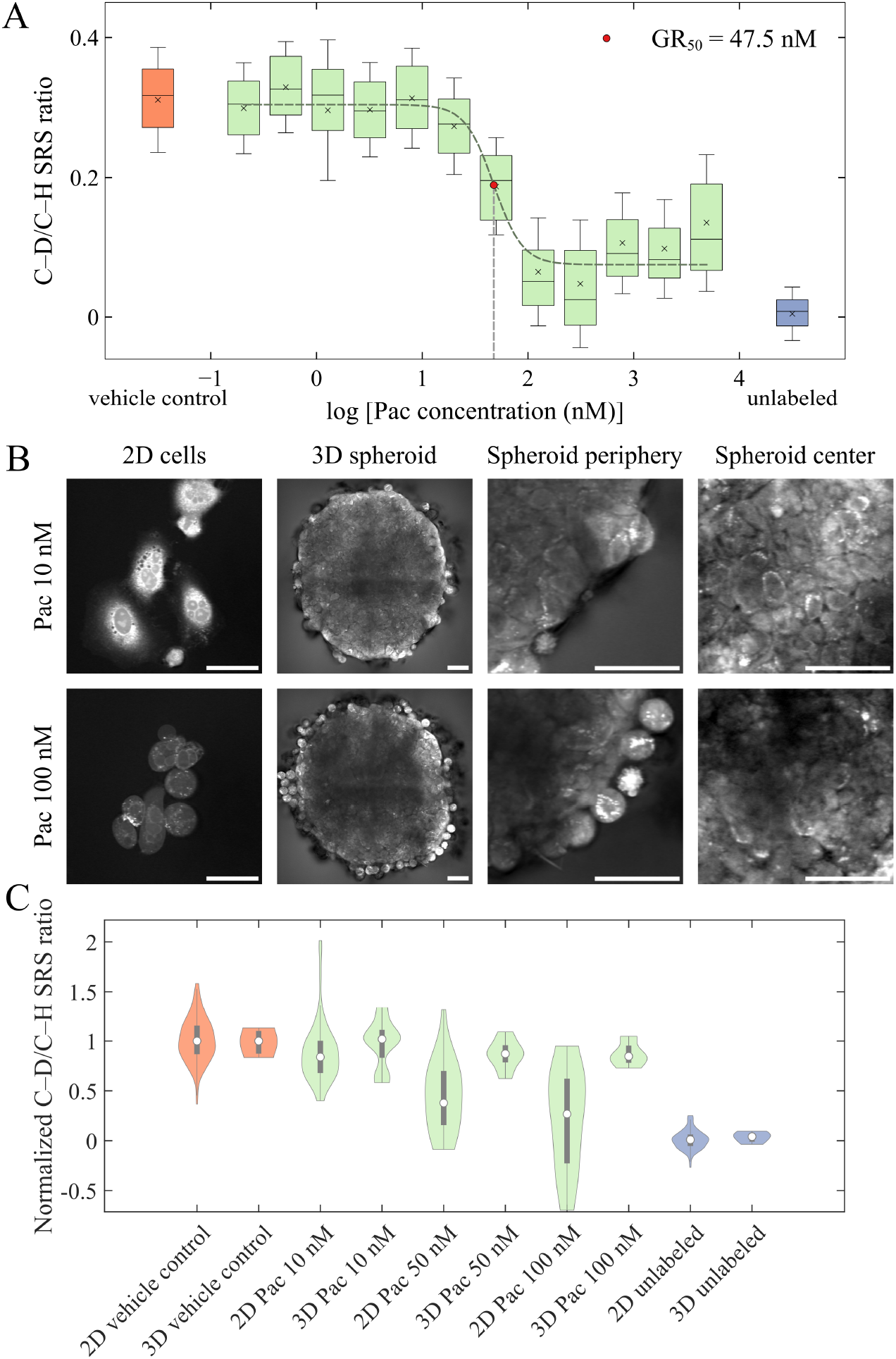
Cellular growth rate inhibition of 2D and 3D A549 cells treated with paclitaxel (Pac). (A)Growth rate response of fixed 2D cells treated with varying concentrations of Pac for 24 hours. (B)Representative C–H SRS images of Pac-treated cells and spheroids. (C) Growth rate inhibition comparison between 2D and 3D cells. Scale bars: 40 μm.

The growth inhibition comparison between 2D and 3D cells shown in **Figure 5C** matches these observations. In both 2D and 3D environments, the cell growth rate decreases as paclitaxel concentration increases. Yet, the decreasing rate, i.e., the growth rate inhibition, is significantly less in 3D, showing the same trend as the lapatinib treatment (**Figure 2C**). Given that the dosed paclitaxel concentrations are below the limit of detection of our SRS method, the drug penetration is not measurable. It is possible that paclitaxel has low penetration through the spheroids, leading to minor growth inhibition.

## Conclusions

Different cell culture architectures and microenvironments lead to distinct cellular responses during drug treatment. Cells cultured in 3D environments generally exhibit higher drug resistance compared to cells in 2D monolayer, but the effect could be highly dependent on drug property and mechanism of action. Understanding such differences is important not only for designing more effective drug screening approaches during drug development but also for optimizing dosage and improving drug treatment in patients. By exploring the spatial features and growth differences between 2D and 3D cells during drug exposure, we can illuminate potential reasons for increased resistance. Our drug imaging efforts complement existing genetic and proteomic techniques that can identify a broad spectrum of genes, proteins, and biochemical pathways contributing to these cellular response differences.

Applying a robust three-band SRS imaging method capable of visualizing and quantifying both intracellular drug uptake and cellular growth response in 2D and 3D models, we performed a quantitative comparative analysis of drug uptake and its growth inhibition across different cell culture systems. Our findings reveal that despite the similar intracellular drug levels in both 2D and 3D A549 cells during treatment with lapatinib, the 3D spheroids exhibit a notably lower impact on their growth, indicating an increased tolerance that likely stems from the complex 3D microenvironment. Our investigations further showed that limited drug penetration contributed to drug gradients in 3D cell cultures. Interestingly, we discovered that even though the inner layer cells in spheroids have less drug exposure and uptake, their growth rates experienced stronger inhibition, likely due to limited nutrient and oxygen availability. We also demonstrated that the presence of ECM can alter the spheroid structure, morphology, and growth and serve as a physical barrier that significantly impedes drug diffusion and uptake. The reduction in drug exposure can then lead to less prominent growth inhibition. These observations highlight how drug efficacy can be significantly modulated by the physical and biochemical environment surrounding the cells. Furthermore, our study extended beyond lapatinib to a chemotherapeutic drug, paclitaxel. Both morphological and quantitative information confirmed that cells in 3D exhibited significantly lower growth inhibition than in 2D under the same treatment. The agreement in spheroids having reduced growth inhibition than monolayer cells between different mechanisms of action drug treatments validated a generally enhanced drug tolerance in the 3D microenvironment.

Our three-band SRS imaging method not only advances our understanding of how biochemical and physical interactions affect drug resistance in 2D and 3D cellular systems but also sets the stage for more sophisticated drug testing with complex 3D culture models such as patient-derived organoids. The compatibility of SRS microscopy with tissue imaging enables the investigation of drug uptake and cell proliferation in patient-derived organoids and tissues. With the incorporation of epi-mode detection, this technique may also allow for in vivo monitoring and measuring drug penetration and cellular response in actual tumors in mice. Additionally, it is important to note that cell growth inhibition does not necessarily correlate with cell death. To enhance the robustness of our findings, our method can be integrated with viability measurements, such as live/dead staining and fluorescence imaging, to provide a more comprehensive profile of cellular responses during treatment in varying microenvironments. Moreover, coupling our results with RNA sequencing could identify specific genetic differences between cells under different culture conditions and drug treatments, thereby refining our understanding of the intricate drug resistance mechanisms. Overall, we anticipate that the continued refinement and application of these methods will yield critical insights necessary for understanding and overcoming drug resistance and improving treatment outcomes in clinical settings.

## Associated Content

### Supporting Information

Intracellular drug uptake measurements with normalization using the 2930 cm^−1^ C–H peak intensity; field-of-view normalization for spheroid imaging (PDF)

## Author Information

### Author Contributions

D.F. and F.X. conceptualized the project. F.X. performed cell and spheroid treatment, SRS imaging, and data analysis. R.S. performed seeding and culturing of both scaffold-free and scaffold-based 3D spheroid. R.O. performed scaffold-free 3D spheroid seeding. K.H. developed and optimized 3D spheroid culture protocols. F.X. and D.F. wrote the manuscript. All authors have given approval to the final version of the manuscript.

