## Supporting Information for "Assessing drug uptake and response differences in 2D and 3D cellular environments using stimulated Raman scattering microscopy"

**Content:**

- **Figure S1:** Intracellular drug uptake measurements with normalization using the 2930 $cm^{-1}$ C–H peak intensity.
- **Figure S2:** Field-of-view (FOV) normalization for spheroid imaging.

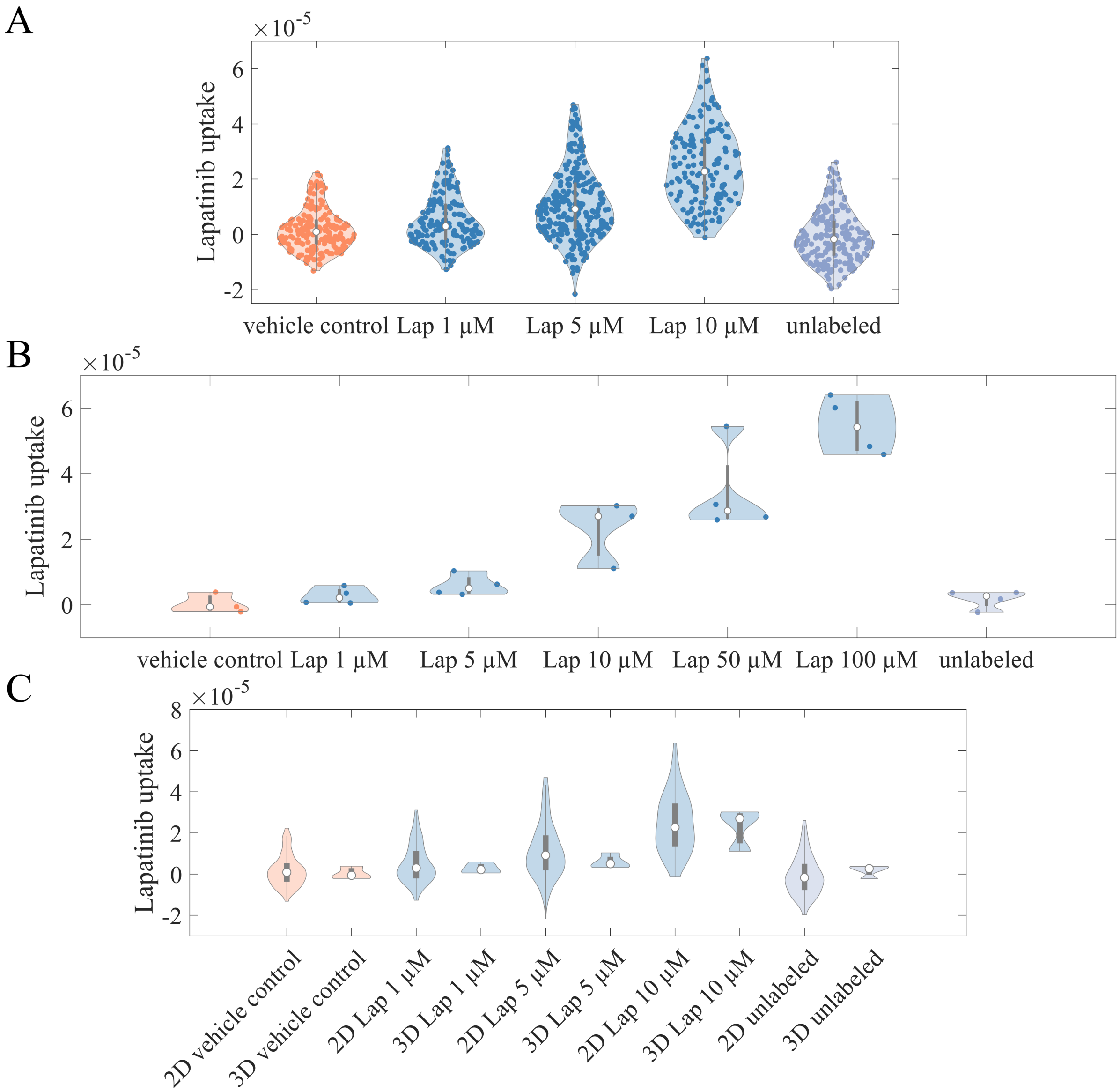


**Figure S1.** Intracellular drug uptake measurements with normalization using the C–H peak intensity at 2930 cm$^{-1}$. (A) 2D A549 cells treated with lapatinib (Lap) of 1–10 µM. (B) 3D A549 spheroids treated with Lap of 1–100 µM. (C) Comparison of 2D and 3D cellular drug uptake.

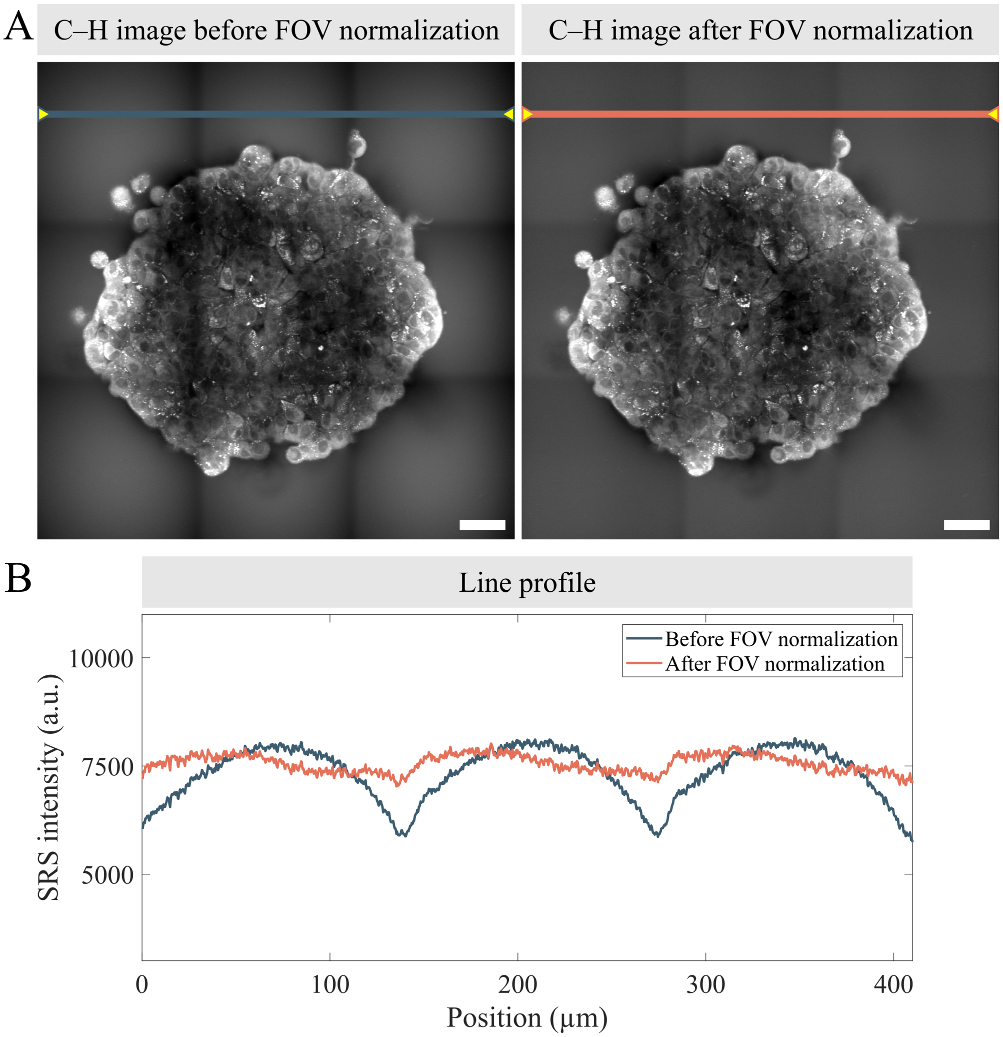


**Figure S2.** Field-of-view (FOV) normalization for spheroid imaging. The natural curvature of each FOV can result in severe stitching artifacts (left in A). We generated FOV normalization masks using cell-free regions and normalized them to have an intensity ranging from 0 to 1. Each SRS image was divided by the FOV mask before stitching. The stitching artifacts were significantly reduced (right in A). (B) Signal intensity profiles before and after FOV normalization. Scale bars: 40 µm.
